# Instantaneous midbrain control of saccade kinematics

**DOI:** 10.1101/305003

**Authors:** Ivan Smalianchuk, Uday Jagadisan, Neeraj J. Gandhi

**Author notes:** Address correspondence to: Neeraj Gandhi.

## Abstract

The ability to interact with our environment requires the brain to transform spatially-represented sensory signals into temporally-encoded motor commands for appropriate control of the relevant effectors. For visually-guided eye movements, or saccades, the superior colliculus (SC) is assumed to be the final stage of spatial representation, and instantaneous control of the movement is achieved through a rate code representation in the lower brain stem. We questioned this dogma and investigated whether SC activity also employs a dynamic rate code, in addition to the spatial representation. Noting that the kinematics of repeated movements exhibits trial-to-trial variability, we regressed instantaneous SC activity with instantaneous eye velocity and found a robust correlation throughout saccade duration. Peak correlation was tightly linked to time of peak velocity, and SC neurons with higher firing rates exhibited stronger correlations. Moreover, the strong correlative relationship was preserved when eye movement profiles were substantially altered by a blink-induced perturbation. These results indicate that the rate code of individual SC neurons can control instantaneous eye velocity, similar to how primary motor cortex controls hand movements, and argue against a serial process for transforming spatially encoded information into a rate code.

## Introduction

Even when we wish to produce the same movement repeatedly, our action exhibits heterogeneity across repetitions. This explains, in part, why intended identical swings of a tennis racket, for example, do not result in identical trajectories of a ball. It is possible the variability could be the result of biological noise in the effectors, although a more likely explanation points to a neural origin (Carmena et al., 2005; Churchland et al., 2006; van Beers, 2007, 2008). While potential neural sources of movement variability have been extensively studied for hand movements (for a review, see (Churchland, 2015)), less is known about their impact on eye movements, particularly the ballistic type known as saccades.

The superior colliculus (SC), a laminar subcortical structure with a topographic organization of the saccade motor map, is a central node in the oculomotor neuraxis (Gandhi and Katnani, 2011; Basso and May, 2017). It is intimately linked to the spatiotemporal transformation, in which visuo-oculomotor signals in the SC conform to a space or place code, while recipient structures in the brainstem exhibit a rate code. In a slight modification to this framework – the so-called dual coding hypothesis (Sparks and Mays, 1990) – saccade amplitude and direction are computed from the locus of population activity in the SC, while movement velocity is a “determinant” of these neurons’ firing rates. The strongest evidence for SC control of saccade velocity comes from causal studies demonstrating that peak eye velocity is correlated with frequency or intensity of electrical microstimulation (Stanford et al., 1996; Katnani and Gandhi, 2012) and that peak velocity is attenuated after inactivation of SC (Sparks et al., 1990). However, these results only address the distribution in static saccade descriptors, falling short of explaining dynamic properties of the movement (e.g. instantaneous velocity). On the other hand, the dynamic vector summation algorithm (Goossens and Van Opstal, 2006) presents a framework in which the SC controls the desired instantaneous displacement of the eye through a series of mini-vectors, but does not directly address the possibility of instantaneous control of ocular kinematics.

In this study, we tested the hypothesis that SC activity dynamically controls the instantaneous velocity of saccades. Time series correlation was first performed on individual trials by regressing the temporal evolution of SC activity with eye velocity within each trial. This analysis, by definition, cannot reveal which epoch(s) of the waveforms contribute most significantly to the correlation. We addressed this latter limitation by correlating the instantaneous neural activity and eye velocity across trials, an ensemble approach that calculates the correlation between firing rate and velocity on individual timepoint basis. We found that instantaneous residual firing rate strongly correlates with instantaneous residual velocity for both within-trial and across-trials analyses. The peak correlation was best aligned with the time of peak eye velocity, and at a population level, the correlation was significant throughout the movement. Neurons with the highest firing rates within individual penetrations displayed the strongest correlation. Finally, the relationship was observed not only for ballistic-like, bell-shaped velocity waveforms of normal saccades but also for profiles altered by blink perturbations. Thus, individual SC neurons exhibit a code that can control instantaneous eye velocity, akin to how primary motor cortex controls hand velocity (Ashe and Georgopoulos, 1994; Reina et al., 2001) Our finding also argues against a serial process for transforming spatially encoded information into a rate code.

## Methods

Two adult rhesus monkeys (*Macaca mulatta*, 1 male and 1 female, ages 8 and 10, respectively) were used for the study. All procedures were approved by the Institutional Animal Care and Use Committee at the University of Pittsburgh and were in compliance with the US Public Health Service policy on the humane care and use of laboratory animals.

Extracellular spiking activity of SC neurons were recorded as head-restrained animals performed a visually-guided, delayed saccade task under real-time control with a LabVIEW-based controller interface (Bryant and Gandhi, 2005). Neural activity was collected with either a multi-contact laminar probe (Alpha Omega; 16 channels, 150 μm inter-contact distance, ~1 MΩ impedance of each contact) or a standard tungsten microelectrode (Microprobes, ~1 MΩ impedance). All electrode penetrations were orthogonal to the SC surface, so that roughly the same motor vector was encoded across the layers. The saccade target was presented either near the center of the neuron’s movement field or at the diametrically opposite location. This study reports analyses from 189 neurons, 145 of which were collected with a laminar probe across 18 sessions, and the remaining 44 neurons were recorded with a single electrode (Jagadisan and Gandhi, 2017). All neurons can be classified as visuomotor or motor neurons according to standard criterion, which we have described previously (Jagadisan and Gandhi, 2016).

Blink perturbation data were only available for the 50 neurons, 44 of which were studied with the single electrode setup and the remaining 7 from a single laminar electrode session. On approximately 15-20% of the trials, an air-puff was delivered to one eye to invoke the trigeminal blink reflex. The puff was timed to induce a blink around the time of saccade onset or even trigger the eye movement prematurely. In this case, blink-triggered saccades provide a valuable control against spurious correlations, as the velocity profile of the saccade is altered compared to that of a normal movement and endpoint accuracy is preserved. Thus, if SC dynamically controls the kinematics of the saccade, the perturbed velocity waveforms should be predicted by the SC activity as well. For full disclosure, the data from these 50 neurons are the same as those reported in a previous publication (Jagadisan & Gandhi 2017). The key distinction is that the previous study assayed SC activity during the saccade preparation phase, and now the focus is on the peri-saccade period.

Eye and eyelid movements were detected using the magnetic search coil method. Spike trains were converted to a spike density waveform by convolution with a 5 ms Gaussian kernel. All movements were aligned on saccade onset and standard velocity criteria were used to detect the onset and offset of normal saccades. For blink-triggered movements, the onset of saccadic component was estimated as the time of a deviation from a spatiotemporal template of a blink related eye movement induced during fixation (Katnani and Gandhi, 2013). The movement profiles were then transformed into radial coordinates in which positive values indicate vectorial velocity towards the movement field and negative values away (Jagadisan and Gandhi, 2017); this differs from the common method in which vectorial velocity representation is always positive and independent of the ideal saccade path towards the target. This distinction is vital, as we were not able to observe the effects described later when using traditional vectorial coordinates. To remove potential confounds of saccade amplitude on correlations between spiking activity and radial velocity, we additionally limited each neuron’s dataset to movements within ±5% of the mean amplitude (range: 8-25 degrees; median: 12 degrees) and, for the analysis comparing normal and blink-perturbed movements, at least 20 trials for each condition. All 189 neurons passed the inclusion criteria, 50 of which also had amplitude-matched blink perturbation trials.

For each neuron we subtracted the session’s mean velocity and spike density waveforms from each trial’s data to obtain the respective residuals. This important step removes spurious correlations from generally similar shapes of SC activity and saccade velocity. For within-trial analysis, a Pearson’s correlation was determined for each trial’s residual velocity and neural activity waveforms. The two residual vectors were shifted relative to each other from 100 *ms* to −100 *ms* in 1 *ms* increments, and the Pearson’s correlation was calculated for each delay (Δ*t*). A zero delay indicates that both vectors are aligned on saccade onset. Negative delays signify instances when the neural activity preceded the velocity, which we refer to as a efferent delay (ED). This analysis was performed for every residual neural activity – velocity waveform pairing for each trial of each cell. For across-trials analysis, we created a vector of activity residuals at time *t* and a corresponding vector of velocity residuals at time *t +* Δ*t*, where the length of the vectors equals the number of trials. The Pearson correlation between these two vectors was determined. We repeated this procedure for every timepoint in the saccade and for every 1 *ms* shift, resulting in a correlation coefficient for each combination of time relative to saccade onset and delay. This technique allowed us to examine the SC effects on eye velocity at every timepoint of the saccade in 1 *ms* resolution. We repeated both correlational analyses on blink-perturbed trials to see if the results persist even when the saccade properties are altered.

To further validate that the influence of the SC on velocity continues throughout the entirety of the saccade, we calculated the duration of significant correlation for each cell. To do this, we determined the confidence threshold by performing the across-trial analysis on shuffled data for each cell one hundred times. Then we determined that the correlation was significant at those points where the real correlation exceeded 2 standard deviations around the mean of the shuffled results. Summing the instances at which correlation was significant gave us the total duration of the correlation.

To determine the significance of the results of within-trial analysis, we randomly paired a residual velocity trace of one trial with residual activity data of another trial from the same session’s data and determined the Pearson correlation for the span of delay values. For across-trials analysis, we randomly shuffled the order of elements in each residual vector (instantaneous across-trial shuffle) before determining the correlation coefficient. The shuffling procedures were repeated 100 times for both types of analyses. Deviations of unshuffled correlation results outside 2 standard deviation bounds generated from shuffled data were deemed statistically significant.

In a separate analysis we looked at the subset of data which was collected using laminar probes. This subset allowed us to examine the effect of depth of SC neurons on our results. The number of channels with neural data ranged from 4 to 16 per session. Since each session did not have enough channels for sufficient statistical power, we de-meaned the data from each session and combined all channels. We then used linear regression on the de-meaned correlation coefficients of these channels and the corresponding FR to establish a trend.

## Results

The relationship between SC activity and saccade dynamics can be intuitively queried by correlating the firing rate with velocity for different transduction delays. Figure 1A plots the results of such within-trial analysis for normal saccades across all 189 neurons in our database. The mean peak Pearson correlation (*r* = 0.179) was observed for an efferent delay (ED) of 13 *ms*, equivalently *Δt* = -13 *ms*. The correlation was significantly different from the pattern observed for shuffled data. This result therefore indicates that the residuals of both SC activity and eye velocity fluctuate around the mean in a coherent fashion and that SC activity can influence saccade velocity.

**Figure 1:**
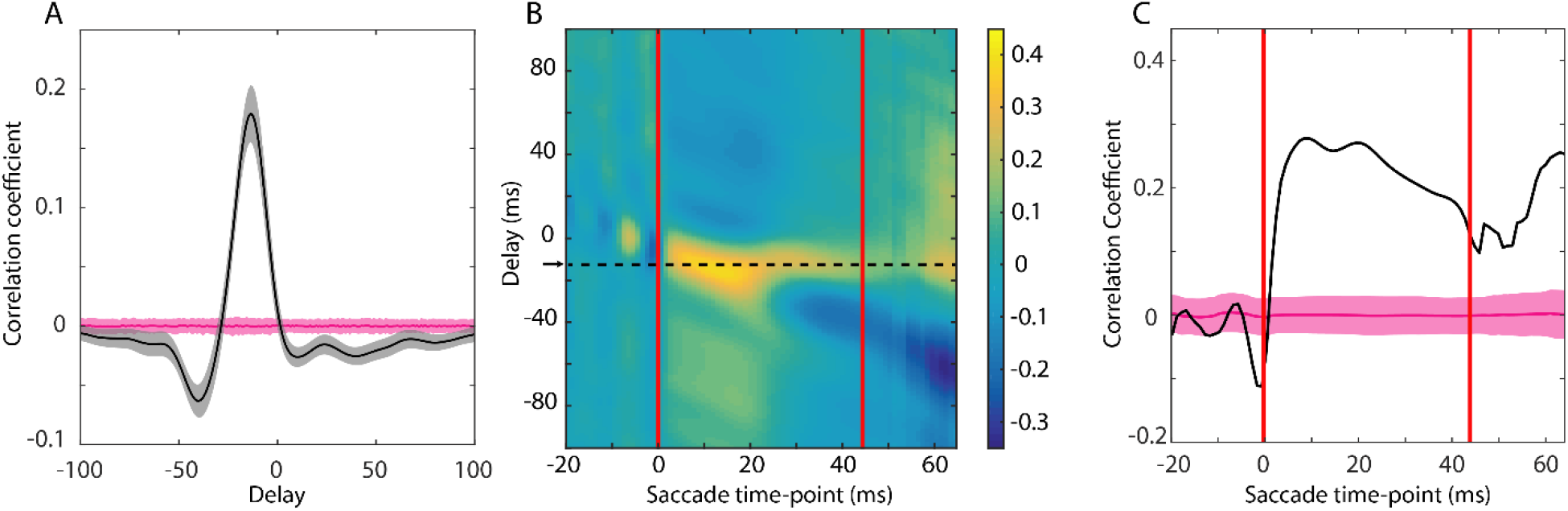
Summary of within– and across-trial correlation analyses for normal saccades. (A) Within-trial correlation analysis. Black line denotes the correlation coefficient between activity and velocity residuals as a function of the temporal shift. The gray outline is two standard errors around the mean patterns from 189 neurons. Pink line and outline represent the mean correlation coefficients and two standard deviations from the mean of the shuffled data. (B) Across-trials correlation analysis. Heatmap of correlation coefficients between SC activity and eye velocity residuals for each timepoint during the saccade (abscissa) and temporal shift between the two residual vectors (ordinate). Arrow and horizontal dashed line mark the efferent delay at which the average correlation was highest (-12ms). Left and right vertical red lines indicate respectively the beginning and the end of the shortest saccade in the dataset. Data past rightmost red bar excludes saccades which have terminated prior to the timepoints on the x-axis. (C) Correlation coefficients as a function of saccade timepoints for the optimal ED shown in (B). Pink line and outline represent the mean and two standard deviations for the across-trials analysis performed on shuffled data.

A major limitation of within-trial analysis is that it offers no information about the correlation at each timepoint within the saccade. The correlation coefficient could peak if any sufficiently long sequence of the activity correlates with the corresponding sequence in velocity. One can imagine that SC could, perhaps, encode the accelerating phase of the eye movement and the deceleration phase could be unaffected by SC activity, and rather be guided by muscle viscoelastic properties. In this case, the correlation would peak at a particular ED because the first half of both signals is correlated, but would provide little evidence to support our hypothesis that SC dynamically influences the entire saccade. We therefore employed across-trials analysis to determine precisely the time-course of correlation between activity and velocity.

Assembled across trials, the residual firing rates were correlated with residual eye velocities separated by a delay. This procedure was repeated for a large range of delays and for all times points of a saccade. Figure 1B shows the correlation coefficients for all combinations of saccade time points and ED values. A horizontal band of high correlation values is noted for the duration of the saccade for an ED of 12 *ms*. The correlation values in this band (Figure 1C) are even higher (peak: *r =* 0.278) than that found in within-trial analysis. Results from the across-trials analysis therefore provide the strongest evidence that SC dynamically influences eye velocity throughout the entire saccade. Additionally, it is prudent to mention that this analysis is identical whether performed on residuals or raw data, thus providing a more direct evidence of correlation as compared to the within-trials analysis.

Next, we explored the temporal characteristics of instantaneous activity-velocity correlations. We found that the time of peak correlation was well aligned with the time of peak velocity, after accounting for the efferent delay, for each neuron (Figure 2A). A paired *t*-test could not reject the null hypothesis that the mean difference was zero (*p* = 0.15). In contrast, the duration of the SC influence over the saccade velocity reveals a flat distribution (Figure 2B), and cells that had a shorter duration of significant correlation tended to have a lower peak correlation (Figure 2C; *r* = 0.53, *p* = 1.99 × 10^-33^, [*f − test, MATLAB regress function*]). The relationship between degree and duration of influence could be explained by cells with higher correlation values having a higher likelihood of rising above the significance level over time. Thus, SC neurons tend to exhibit most influence over eye kinematics around the peak of the saccade velocity profile, and those cells which showed a higher peak correlation continued to influence eye velocity well past the peak, to saccade completion.

**Figure 2:**
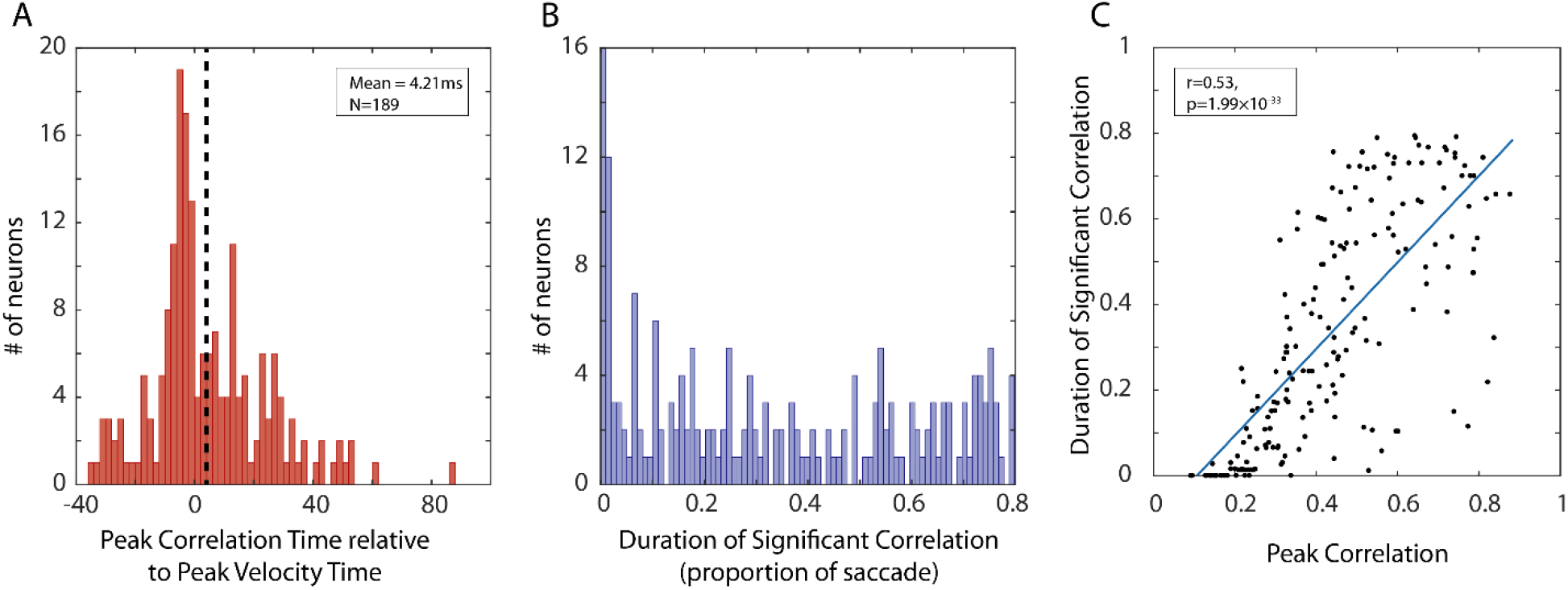
Temporal characteristics of activity-velocity correlation. (A) Histogram of average peak correlation time relative to average peak velocity time for each neuron. The count on y-axis indicates the number of neurons. (B) Histogram of cumulative duration (as proportion of total saccade length) for which the correlation remained above significance level. (C) Relationship between peak correlation and the duration of the correlation. Each point indicates one neuron. Blue line is the best fit line to the data.

We then studied which properties of the neuronal population contributed to significant correlations. When examining the entire population of cells, simple linear models found no relationship between a cell’s peak correlation and its peak firing rate or its location along the rostral-caudal extent of the SC (*p* = 0.088, *f − test*). However, when we isolated only the cells from a single laminar recording, we observed an increasing trend between the cells FR and its peak correlation (Figure 3A). When data from the laminar recordings was pooled by subtracting the average peak firing rate and correlation measures of each session, a statistically significant linear relationship was observed (Figure 3B, *p* = 7.4 × 10^-6^, *f − test*). This suggests that there is a strong relationship between firing rate and instantaneous velocity within individual columns of the SC.

**Figure 3:**
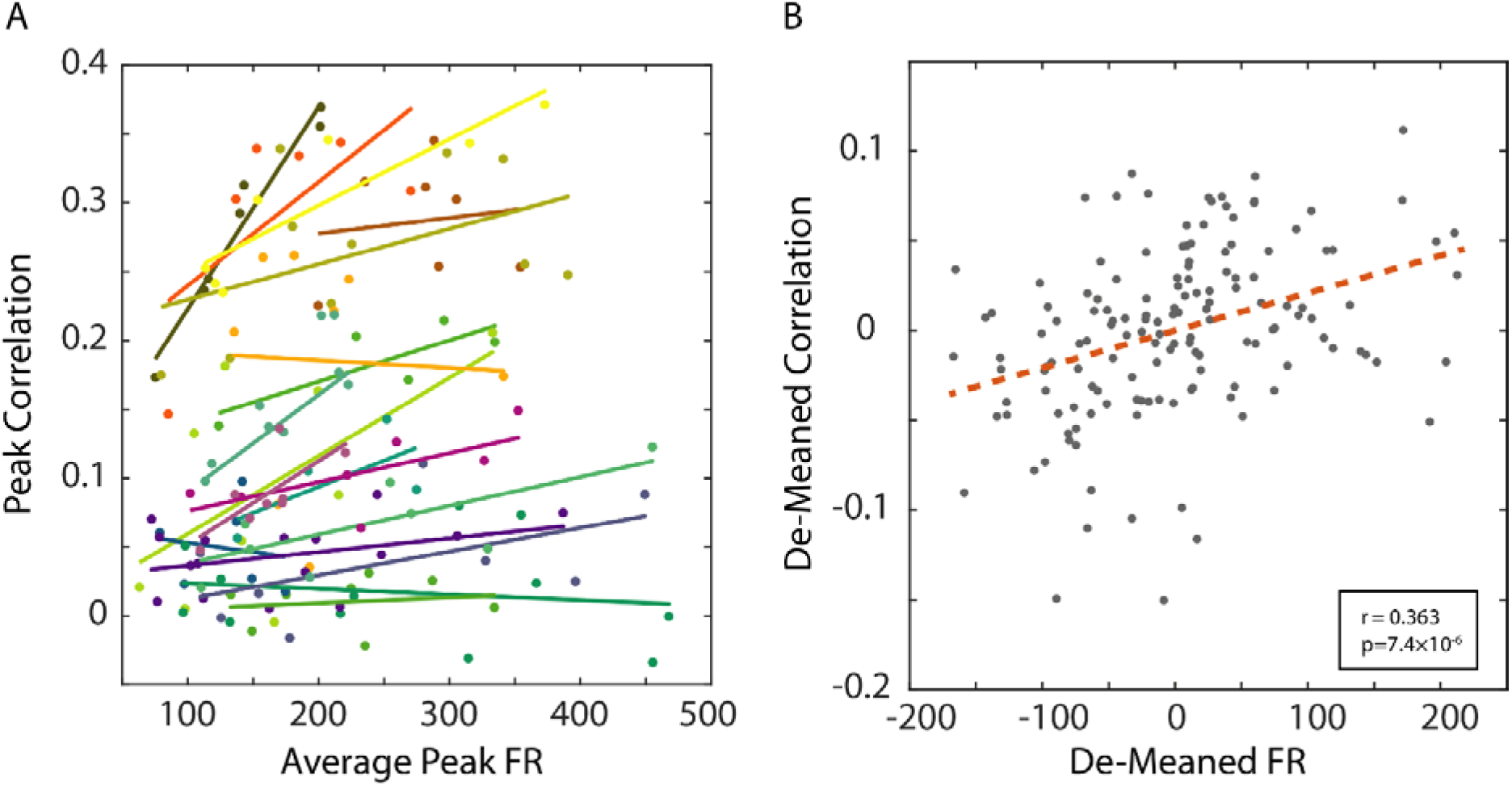
Analysis of data recorded from laminar probes. (A) Peak correlation of each SC neuron is plotted against the average peak firing rate of that neuron. Neurons recorded in the same penetration are plotted as the same color, each color represents data from different sessions. The best fit line to each session’s data is shown in the matching color. (B) Data from (A) de-meaned and pooled across sessions. Each de-meaned value is obtained after subtracting the respective average across all neurons in its track. The dashed red line is the best fit line.

To assess the robustness of the influence of SC activity over instantaneous eye velocity we turned to the 50 neurons for which we also had blink-perturbation data. Such saccades do not exhibit the stereotypical, bell-shaped profile and therefore offer an opportunity to assess if the correlation persists even in the presence of perturbation. The top row of Figure 4 displays the within– and across-trial analyses for normal trials in the 50 neurons. The general trend is preserved, even with the majority of data removed. The best EDs for the normal, unperturbed data were 11 *ms* and 12 *ms* and the peak correlations were *r* = 0.267 and *r* = 0.441 for within– and across-trial analyses, respectively. For the blink-perturbed data from the same neurons, the peak correlations were *r* = 0.210 and *r* = 0.386 for both within– and across-trial analyses, respectively; and the ED was 13 for both. Correlative relationship was largely intact despite the blink-induced perturbation.

**Figure 4:**
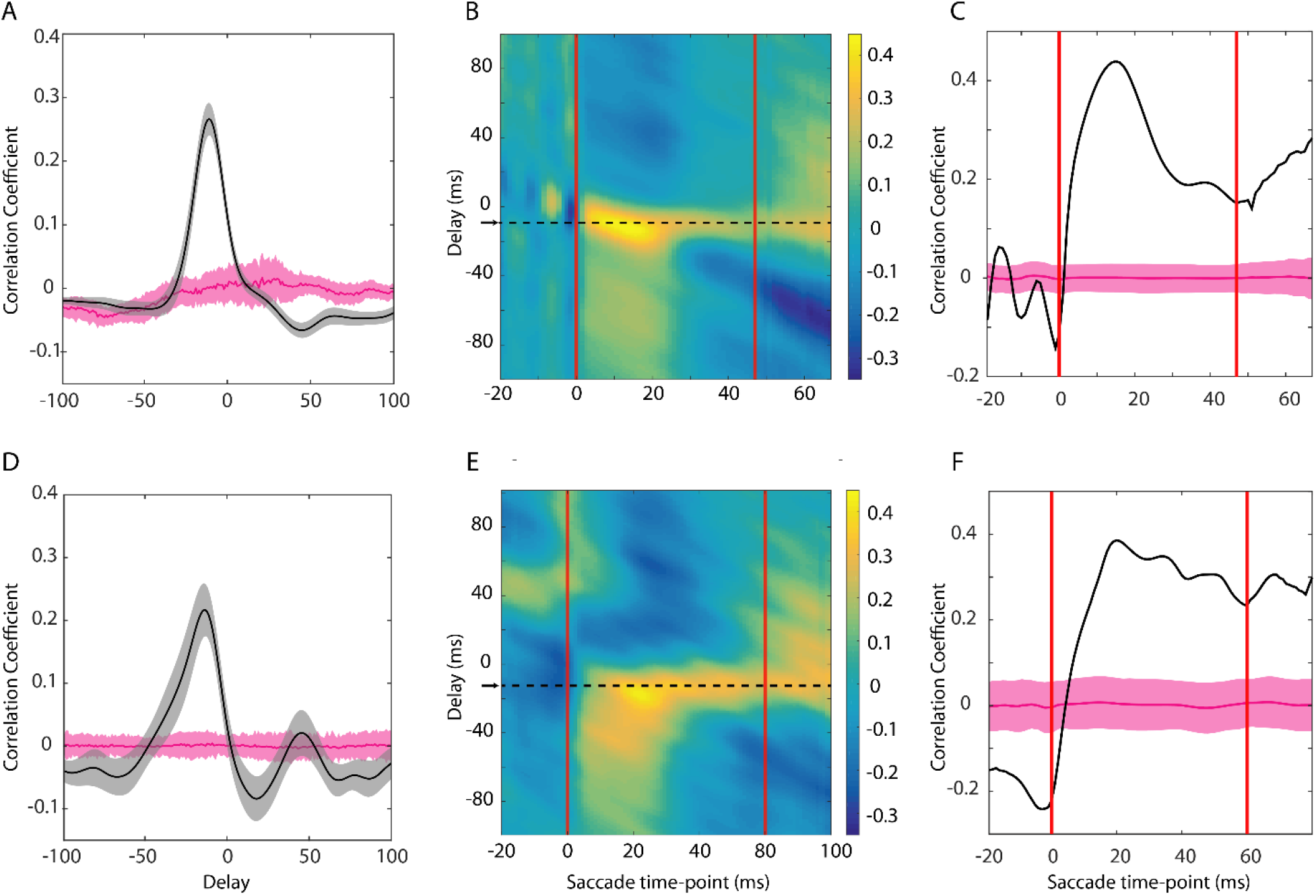
Comparison of correlation analyses for blink-perturbed and normal saccades. Within-trial correlation between activity and velocity residuals for (A) normal and (D) blink-perturbed saccades available for 50 of 189 neurons. The heatmaps of correlation coefficients obtained from across-trials analysis for (B) normal and (E) blink-perturbed movements. Correlation coefficients as a function of saccade timepoints for the optimal ED for (C) normal and (F) blink-perturbed saccades. The plots follow the same conventions used in Figure 1.

To further characterize the impact of blink perturbation, we computed for each neuron the linear relationship between residual activity and velocity distributions across the duration of the movement.

Assuming the data contained *n trials* and the average duration of the amplitude-matched saccades is *d ms*, a single regression was performed across *nd* points. The activity data were time shifted relative to the velocity distribution to account for the neuron’s ED. This was done separately for the normal and blink-perturbed data from each of the 50 neurons. Figure 5A shows a scatter plot of the regression slopes for each neuron in the two conditions. Overall, the slopes tended to be greater during the blink condition (paired t-test p=0.0013). Moreover, we found a strong relationship between each neuron’s regression slope and the goodness of fit in both normal (*r*=0.72, p=2.37×10^-7^) and perturbation (*r*=0.35, p=0.02) conditions (Figure 5B).

**Figure 5:**
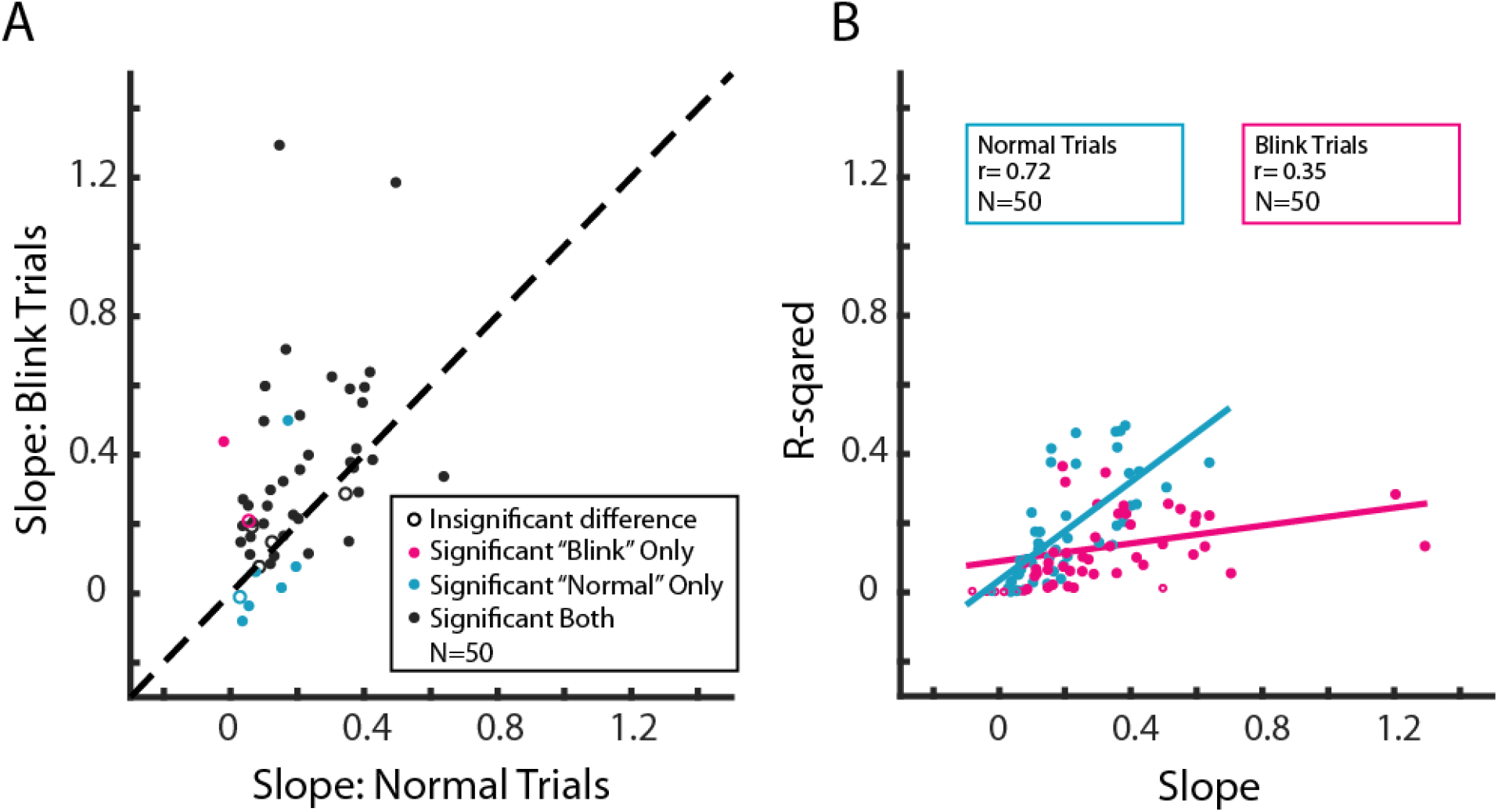
Linear regression features between SC activity and eye velocity. (A) A pairwise comparison of the regression slopes obtained for normal (x-axis) and blink-perturbed (y-axis) conditions for each neuron. Every neuron had a statistically significant slope for at least one of the two conditions. Cyan: normal trials only; magenta: blink perturbation trials only; black: both types of trials. The dashed line represents the unity relationship. (B) Relationship between slope and r^2^ values. Cyan: normal trials. Magenta: blink perturbation trials.

## Discussion

We revealed an intuitive yet, to our knowledge, unreported phenomenon on how SC activity exerts instantaneous control over saccadic eye movements. We found a strong correlation between the motor burst of SC neurons and eye velocity for an efferent delay of approximately 12 *ms*. The correlation was noted for both within-trial and across-trials analyses. The latter approach, in particular, demonstrated that the correlation remained high for the duration of saccade, lending support for SC control of instantaneous eye speed and thereby modifying the notion of spatiotemporal transformation. In addition, this approach revealed a robust distributed population coding scheme reminiscent of a synfire chain (Diesmann et al., 1999; Shmiel et al., 2006), wherein individual neurons exert influence over saccade dynamics sequentially at different times, collectively spanning the duration of the saccade. Comparable correlation structure and ED were also observed for blink-perturbed movements, whose velocity profiles deviate significantly from the stereotypical bell-shaped waveforms (Goossens and Van Opstal, 2000; Gandhi and Bonadonna, 2005), although it was particularly important to project the velocity vector onto the preferred vector of the SC neuron (Jagadisan and Gandhi, 2017). We also learned that, within individual collicular columns, cells with higher FR tend to exhibit a stronger correlation with velocity, suggesting a relationship based on the depth of each cell as well as its spiking properties. Finally, we uncovered the interesting feature that regressions with the largest correlations also had the biggest slopes, implying that eye velocity is more sensitive to firing rates of the more reliable neurons.

Previous studies have addressed relationships between neural activity and movement parameters in several ways. For SC control of saccades, the focus has been on static parameters. For example, weaker bursts of activity produce saccades with lower peak velocity (Edelman and Goldberg, 2001), peak velocity is correlated with frequency or intensity of microstimulation (Stanford et al., 1996; Katnani and Gandhi, 2012), and peak velocity is attenuated after inactivation of SC (Sparks et al., 1990). Modest correlations with SC activity have also been reported for head movements (Walton et al., 2007; Rezvani and Corneil, 2008) and electromyographic activity in proximal limb muscles (Stuphorn et al., 1999). Instantaneous control of saccades has been studied or discussed for neurons downstream of the SC, in the oculomotor brainstem (Cullen and Guitton, 1997; Sylvestre and Cullen, 1999; Sparks and Gandhi, 2003). Such analyses are also more common in the skeletomotor system, with significant correlations identified between neural activity waveforms in cortical areas and hand velocity profiles (Ashe and Georgopoulos, 1994; Reina et al., 2001). Trial-to-trial variability in eye and hand velocity has also been attributed to variability in neural activity occurring earlier in the trial, for instance during sensory input (Osborne et al., 2005; Huang and Lisberger, 2009) and motor preparation (Churchland et al., 2006; Jagadisan and Gandhi, 2017), but this perspective precludes insight into direct dynamic control during ongoing movements.

We believe our study is the first to quantify correlations informative of instantaneous control of saccades by SC neurons. This finding does not conform readily to the standard role of SC in saccade generation, that the spatial distribution of population activity in the deeper layers determines the saccade vector and that downstream structures generate the firing patterns that reflect the velocity profile of the eye movement. The result that SC activity does influence peak saccade velocity (cited above) somewhat aligns to a modified framework, the dual coding hypothesis (Sparks and Mays, 1990), in which the level of SC activity acts as a gain factor on the brainstem burst generator (Nichols and Sparks, 1996). However, influencing saccade speed through a global gain is not equivalent to imposing instantaneous control. In contrast, a dynamic vector summation algorithm (Goossens and Van Opstal, 2006; Goossens and van Opstal, 2012) offers a more compelling framework in which the SC controls the *desired* instantaneous displacement of the eye. Crucially, this model abstains from making direct statements about instantaneous velocity control. It is possible this is because SC neurons do not have a relationship between the average firing rate and the velocity of the saccade they code for. In other words, low firing cells could code for high-velocity saccades, and high-firing cells could code for low-velocity saccades, depending on where they reside in the collicular map. Our analysis circumvents this limitation by examining residuals. We suggest that the static parameters of the saccade are controlled by traditionally understood methods (e.g. dual coding hypothesis) while deviations from the optimal velocity profile are influenced by dynamic fluctuations of neural activity residuals. This proposed framework is capable of functioning in tandem with the aforementioned dynamic vector summation model, as long as the model accounts for the fluctuations in the residual activity of each cell.

The efferent delay is indicative of the transduction time of neural signals from the SC to the extraocular muscles. Studies of SC stimulation have established a 25 − 30 *ms* latency for movement initiation (Stanford et al., 1996; Katnani and Gandhi, 2012) and a shorter 10 − 12 *ms* delay for perturbing an ongoing movement (Munoz and Wurtz, 1993; Miyashita and Hikosaka, 1996; Gandhi and Keller, 1999). The ED that yielded the strongest correlation from our neural recording data was approximately 12 *ms*. Crucially, it remained relatively constant throughout the movement, although we did observe a broader range of ED values with higher correlation coefficients around saccade onset (Figure 1B). While the ED values may seem different for microstimulation and recordings studies, a direct comparison should be avoided because underlying network level processes associated with movement preparation, which are implicitly incorporated in neural activity, may be different when microstimulation is used to trigger a movement.

We are intrigued by the observation that peak correlation between activity and velocity increased as a function of peak firing rate, but only for neurons within a SC “column” and not across the entire population (Figure 3A). Analyses from separate, unpublished work in our laboratory indicate that peak firing rate of the motor burst changes nonlinearly with depth, reaching a maximum in the intermediate layers and decreasing gradually for dorsal and ventral locations. Neurons with the highest firing rates resemble the classical saccade-related burst neurons, some with buildup activity, and are likely those that project to the gaze centers in the brainstem (May, 2005; Rodgers et al., 2006).

Finally, we found it interesting that the regression slopes tended to be higher for blink-perturbation trials than for unperturbed movements (Figure 5A). We speculate that reacceleration of saccades in the blink-perturbation condition (Goossens and Van Opstal, 2000; Gandhi and Bonadonna, 2005) produces residuals that likely increase the regression slope. However, we also note that the variance accounted for (r^2^) by these fits were lower for the blink condition (Figure 5B). This suggests that although SC activity is updated to reflect the change in velocity, this compensation is likely not complete and that additional temporal control signals are added downstream in the brainstem burst generator.

In summary, we present here compelling evidence that the SC has dynamic influence over every aspect of the saccade. The result impacts the notion of spatiotemporal transformation, which is thought to be a serial process of first encoding the movement in a retinotopic reference frame (place code) and then transforming it into a rate code to control its dynamics (Groh, 2001). The SC is considered the last stage of the spatial representation and gaze centers in the lower brainstem employ the temporal algorithm. Our analyses show a clear role of the SC in also exerting temporal control over the duration of the movement, casting a shadow on a simplistic and serial sensorimotor transformation framework. In doing so, the result aligns well with observations from skeletomotor research, where it was demonstrated decades ago that neurons in the cortex encode velocity of hand movement (Ashe and Georgopoulos, 1994; Reina et al., 2001).

## Acknowledgements

Funding for this reseearch was provided by the following grants: NIH T32 DC011499, R01 EY022854, R01 EY024831, and F31 EY027688.

